# Surface motility favors co-dependent interaction between *Pseudomonas aeruginosa* and *Burkholderia cenocepacia*

**DOI:** 10.1101/2022.03.16.484684

**Authors:** Charles Morin, May Landry, Marie-Christine Groleau, Eric Déziel

## Abstract

Interactions between different bacterial species shape bacterial communities and their environments. The opportunistic pathogens *Pseudomonas aeruginosa* and *Burkholderia cenocepacia* both can colonize the lungs of individuals affected by cystic fibrosis. Using the social surface behavior called swarming motility as a study model of interactions, we noticed intricate interactions between *B. cenocepacia* K56-2 and *P. aeruginosa* PA14. While strain K56-2 does not swarm under *P. aeruginosa* favorable swarming conditions, co-inoculation with a non-motile PA14 flagellum-less Δ*fliC* mutant restored spreading for both strains. We show that *P. aeruginosa* provides the wetting agent rhamnolipids allowing K56-2 to perform swarming motility, while aflagellated PA14 seems able to «hitchhike» along with K56-2 cells in the swarming colony.

**Importance:** *Pseudomonas aeruginosa* and *Burkholderia cenocepacia* are important opportunistic pathogens often found together in the airways of persons with cystic fibrosis. Laboratory co-culture of both species often ends with one taking over the other. We used a surface motility assay to study the social interactions between population of these bacterial species. Under our conditions, *B. cenocepacia* cannot swarm without supplementation of the wetting agent produced by *P. aeruginosa*. In a mixed colony of both species, an aflagellated mutant of *P. aeruginosa* provides the necessary wetting agent to *B. cenocepacia*, allowing both bacteria to swarm and colonize a surface. We highlight this peculiar interaction where both bacteria set aside their antagonistic tendencies to cooperate.

## 1 Introduction

Studying polymicrobial communities and their complexity is a priority question in microbiology (1). Social interactions between multiple bacterial species in a shared environment can be either collaborative or competitive. These interactions can have major effects on the community, notably on growth and survival (2, 3). Polymicrobial interactions can be responsible for the increase in antibiotic resistance and the development of persistent infections (4). The airways of people with cystic fibrosis (CF) constitutes a very diverse polymicrobial environment. CF is an hereditary autosomal recessive disease where affected individuals become more likely to develop chronic endobronchial infections (5). The diverse CF airways ecosystem is composed of bacteria, viruses, and fungi (6, 7). Among the bacterial species associated with CF colonization in patients are *Pseudomonas aeruginosa*, and members of the *Burkholderia cepacia* complex (5, 8–10).

*P. aeruginosa* is a Gram-negative opportunistic pathogen infecting immunocompromised individuals and patients with defective barrier defenses (11). When chronically infecting in CF lungs, it induces a lethal decline in respiratory function (5, 12). The polar flagellum of *P. aeruginosa* promotes different motilities such as swimming and swarming (13, 14). Swarming motility consists of a rapid coordinated group movement on a semi-solid surface (typically 0.5 % agar) and requires a functional flagellum and the production of a surfactant acting as a wetting agent (13, 15, 16). The wetting agent produced by *P. aeruginosa* to achieve swarming motility consists of rhamnolipids, extracellular metabolites whose synthesis is regulated by quorum sensing (14, 17–19).

*Burkholderia* species can be isolated from diverse environments such as plant rhizosphere, water, and soil in general (20, 21). *Burkholderia cenocepacia* is a Gram-negative flagellated member of the ever-expanding *Burkholderia cepacia* complex which regroups more than 20 species (22, 23). *B. cenocepacia* is also able to swarm on semi-solid media (24, 25).

Interactions between both opportunistic pathogens have been investigated: *B. cenocepacia* stimulates biofilm production by *P. aeruginosa* by intensifying its biomass (8). When both species are co-cultured *in vitro, P. aeruginosa* can outcompete or inhibit *B. cenocepacia* due to the release of distinct toxic compounds such as hydrogen cyanide (8, 26, 27). Rhamnolipids from *P. aeruginosa* also affect the colony shape of *B. cenocepacia* and promote its swarming (25).

Here, to gain knowledge on collaboration and competition between saprophytes that are also opportunistic pathogens, which could be relevant for understanding progression of infectious colonization and multi-species pathogenesis, we investigated interactions between *P. aeruginosa* PA14 and *B. cenocepacia* K56-2 in a swarming colony using surface motility as a behavioral study system.

## 2 Results

### 2.1 *B. cenocepacia* K56-2 can exploit rhamnolipids produced by *P. aeruginosa* PA14 to achieve a swarming behaviour

*P*. aeruginosa PA14 and *B. cenocepacia* K56-2 are both motile bacteria capable of performing swarming motility on semi-solid media. They both possess two essential characteristics to swarm: a functional flagellum and the production of a surfactant. But while swarming motility of K56-2 is observed on nutrient broth plus 0.5 % glucose media (25), it is unable to spread on M9DCAA swarming media—which is used for *P. aeruginosa*—and rather remains at the inoculation site **(Figure 1)**. *B. cenocepacia* possesses a *rhl* operon homologous to the one from *B. thailandensis* and *B. glumae*, which directs the production of rhamnolipids in these two species (28–30). A mutant of *B. cenocepacia* K56-2 in the first gene of the operon causes a loss of swarming motility on swarming-permissive nutrient media **(Supplemental Fig. S1)**. It is thus very likely that this operon is responsible for the production of a surfactant although we have not been able to identify its structure yet. Since it appears that production of the unidentified surfactant of K56-2 is deficient under *P. aeruginosa* swarming conditions **(Fig. S1)**, we aimed to complement the swarming defect of K56-2 by adding exogenous rhamnolipids or co-inoculating with rhamnolipid-producing *P. aeruginosa*. Adding rhamnolipids on top of the semi-solid gel prior to the inoculation allowed K56-2 to spread on the surface **(Fig. 1B)**.

**Figure 1.**
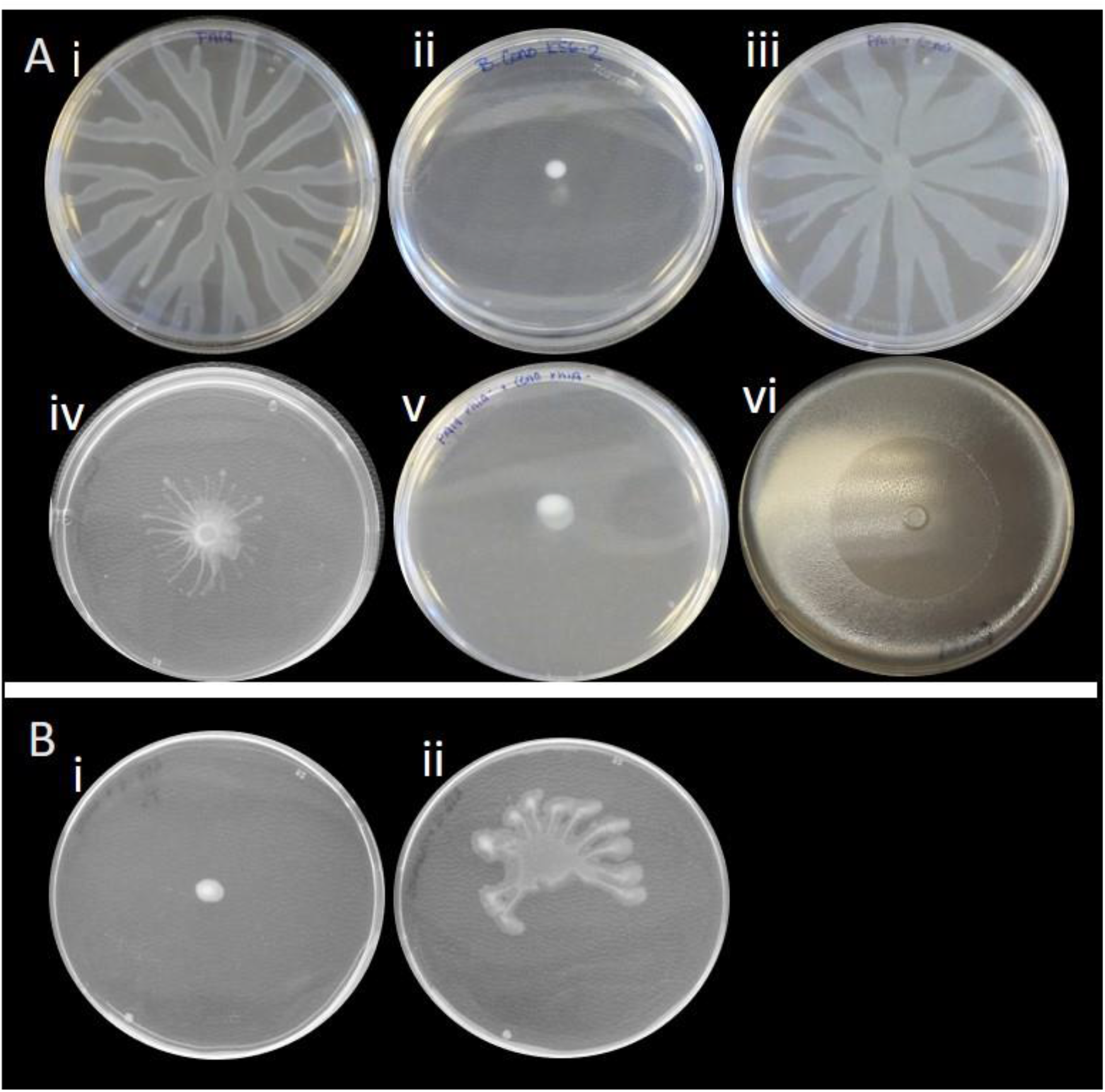
Co-swarming of *P. aeruginosa* PA14 and *B. cenocepacia* K56-2. (**A)** Swarming on M9DCAA semi-solid medium at 30°C: i) *P. aeruginosa* PA14 alone. ii) *B. cenocepacia* K56-2 alone. iii) Mixed 1:1 population of *P. aeruginosa* PA14 and *B. cenocepacia* K56-2. iv) PA14 Δ*fliC* with K56-2. v) PA14 *rhlA*-with K56-2. vi) Oil vaporization highlights the biosurfactant diffusion zone produced by non-motile PA14 Δ*fliC* mutant. (**B)** K56-2 on M9DCAA semi-solid medium supplemented with: i) methanol or ii) with rhamnolipids (dissolved in methanol).

Since strain K56-2 can use rhamnolipids from PA14 to swarm by itself, we hypothesized that K56-2 would swarm alongside PA14 on M9DCAA swarming media. We thus conducted co-swarming assays by mixing K56-2 with PA14. When K56-2 is mixed with PA14, it causes a noticeable alteration in the swarm pattern compared to wild type PA14 alone **(Fig. 1A)**. However, when a rhamnolipid-deficient *rhlA*-mutant of *P. aeruginosa* is instead co-inoculated with K56-2, no swarming is observed, as would be expected if the production of a wetting agent is a requirement for swarming motility under these conditions.

### 2.2 Non-flagellated PA14 Δ*fliC* mutant can spread in a swarm when co-inoculated with K56-2

Providing exogenous rhamnolipids induces the swarming of K56-2; adding K56-2 to a PA14 changes the swarming pattern of the colony. This suggests co-migration between both species. Since a flagellum-null mutant (Δ*fliC*) of *P. aeruginosa* produces its wetting agents rhamnolipids while still being unable to swarm alone **(Fig. 1A vi)**, we reasoned that mixing PA14 Δ*fliC* with K56-2 would allow the latter to swarm. Indeed, swarming is observed when this mutant is co-inoculated with K56-2, suggesting that the Δ*fliC* strain provides rhamnolipids necessary for K56-2 to swarm (Figure 1A iv).

To better understand how two strains unable to swarm when grown separately can produce a swarming colony when cultured together, we labelled PA14 and its Δ*fliC* mutant with the mCherry fluorescent protein. Unexpectedly, when the PA14 Δ*fliC* mutant is co-inoculated with K56-2, it can spread throughout the swarm tendrils, although it remains absent from the tendril tips **(Fig. 2B)**, in contrast with wildtype PA14 in co-swarming **(Fig. 2A)**. This was confirmed with higher magnification using stereomicroscopy **(Figs 2C-F)**. We failed to precisely image K56-2 tagged with GFP due to high background green fluorescence from PA14. To circumvent this limitation, we used CLSM to confirm the position of both bacterial species in the same tendril. **Figure 3** shows the localization of both species in a tendril during co-swarming. Under these conditions, PA14 is prevailing at the tendril tips and leads the way while K56-2 appears to border the tendrils as if being pushed by PA14. When the non-flagellated PA14 mutant co-swarms with K56-2, the latter is more abundant at the tips and leads the tendrils, while PA14 Δ*fliC* trails behind. Both species also appear to be segregated into two distinct populations, with little overlap between them. It is unclear how PA14 follows K56-2 in the swarm without its flagellum.

**Figure 2.**
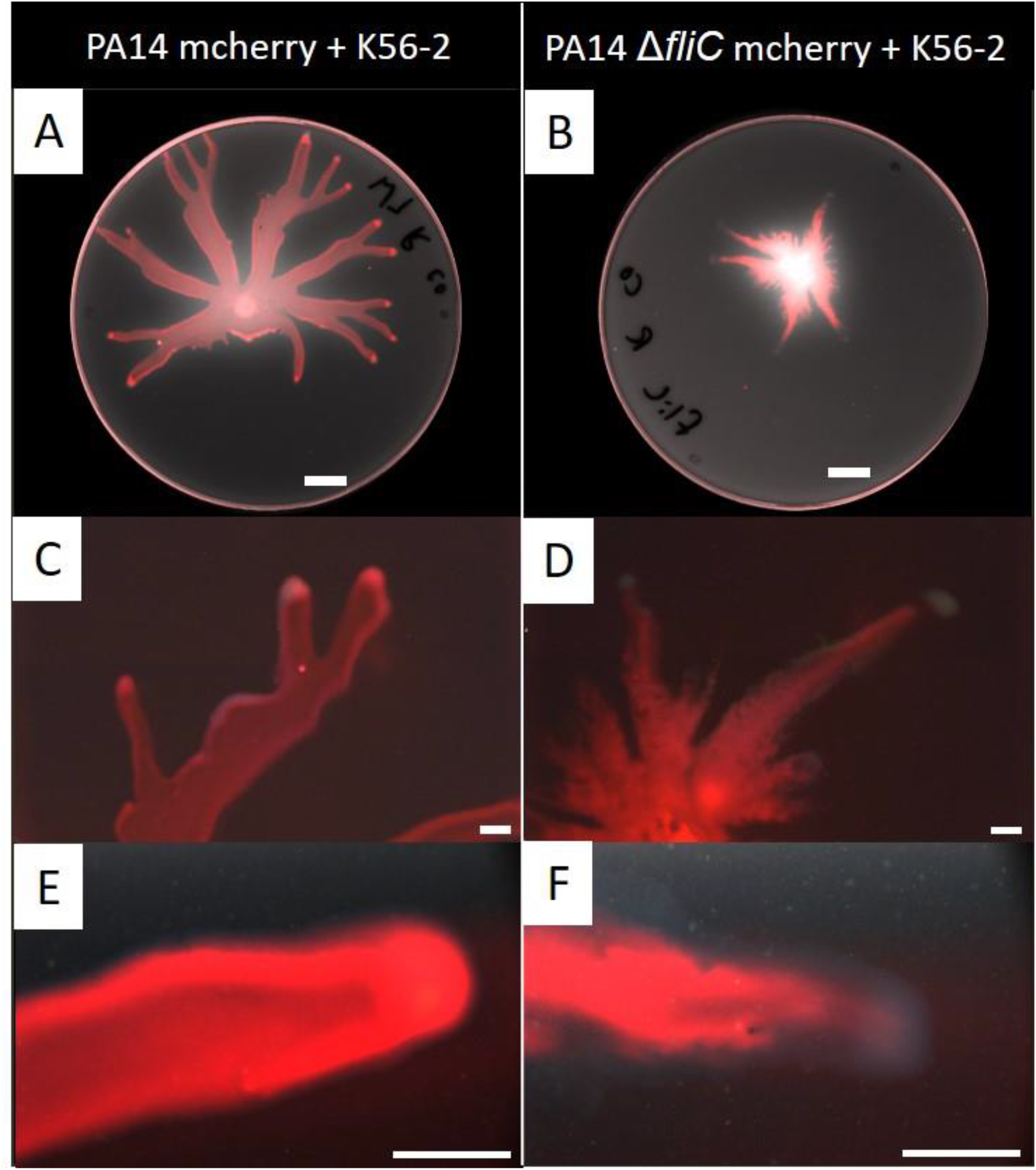
Localization of fluorescently labelled *P. aeruginosa* PA14 in a co-swarm with *B. cenocepacia* K56-2. **LEFT** PA14 (red) with K56-2 (1:1 ratio). **RIGHT** PA14 Δ*fliC* (red) with K56-2 (1:1 ratio). **(A-B)** Images taken with Typhoon FLA9000 (white scale bar = 1 cm). Red color shows mCherry-labelled PA14 against the autofluorescence measured in the green channel shown as grayscale. (**C-D)** and **(E-F)** Images taken with an Olympus SZX16 stereomicroscope at 3.5X and 14X magnification, respectively (white scale bar = 2 mm). Red is the mCherry-labelled PA14 against the whole colony visualized in darkfield.

**Figure 3.**
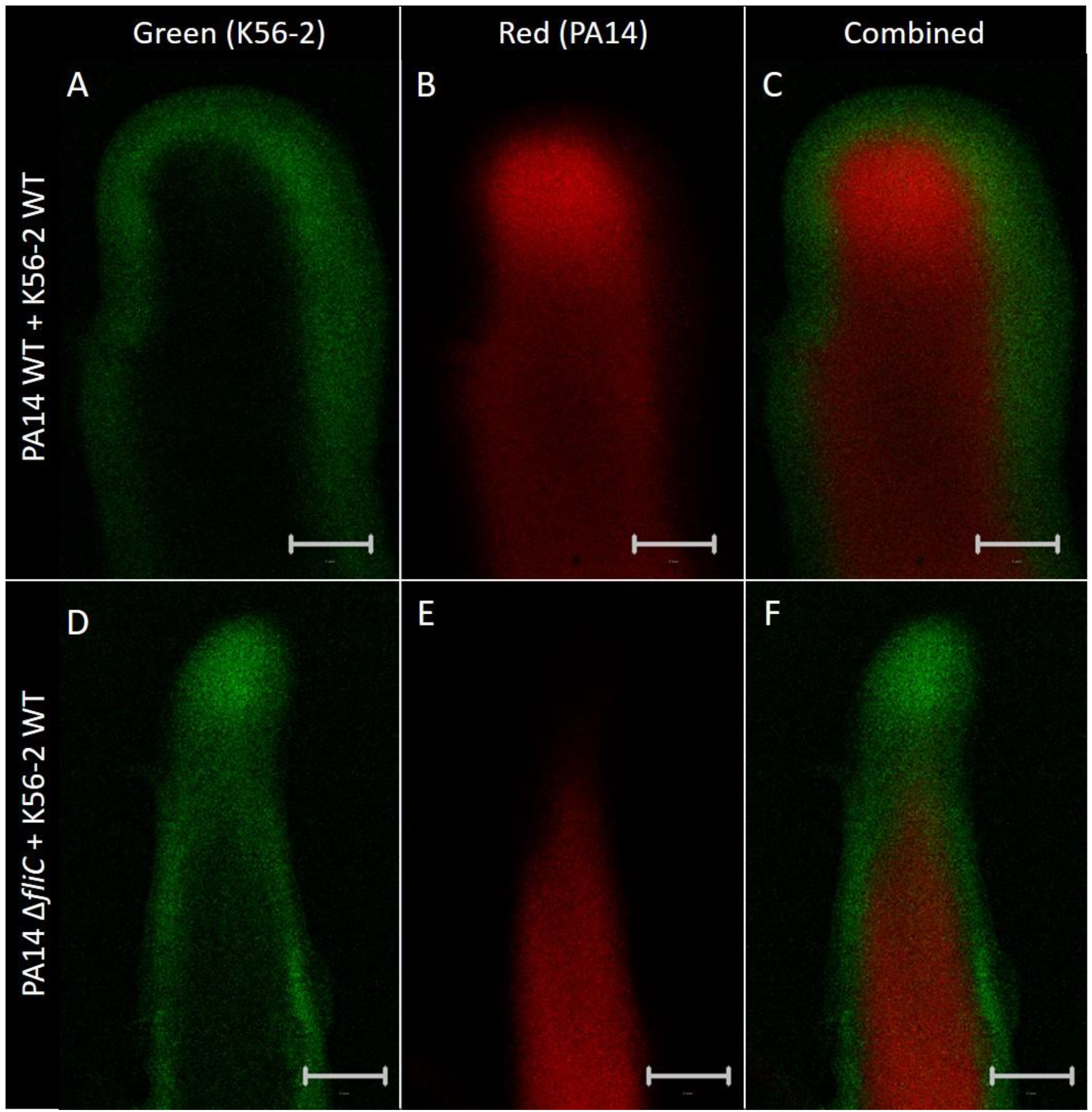
CLSM imaging of *P. aeruginosa* and *B. cenocepacia* co-swarming. Images acquired after overnight growth on semi-solid M9DCAA medium, from a tendril-bearing agar pad. PA14 is labelled with mCherry and K56-2 with eGFP (green) **TOP** PA14 and K56-2. **BOTTOM** PA14 Δ*fliC* with K56-2. Scale bar = 1 mm.

### 2.3 Investigating the appendages required for interactions between *P. aeruginosa* and *B. cenocepacia* in a swarming colony

To determine which *P. aeruginosa* cellular appendage plays a role in the interaction between PA14 and K56-2, we assessed the co-swarming potential of the PA14 Δ*fliC* mutant lacking either the type IV pili (T4P), the CupA fimbriae and/or the Tad pili. These appendages were selected based on their general importance in surface motility and attachment (14, 25). The absence of any of these appendages did not cause a loss of bacterial movement of the PA14 Δ*fliC* mutant in co-swarming with K56-2, although the lack of T4P altered the shape of the swarming pattern **(Fig. 4)**. These results could not support a model of interaction through a specific appendage. To investigate this hypothesis further, we assessed the ability of *P. aeruginosa* swarming cells to carry inert fluorescent polystyrene beads. While PA14 alone or combined with K56-2 were able to move the beads along, a mixture of PA14 Δ*fliC* and K56-2 was unable to do so **(Fig. 5)**.

**Figure 4.**
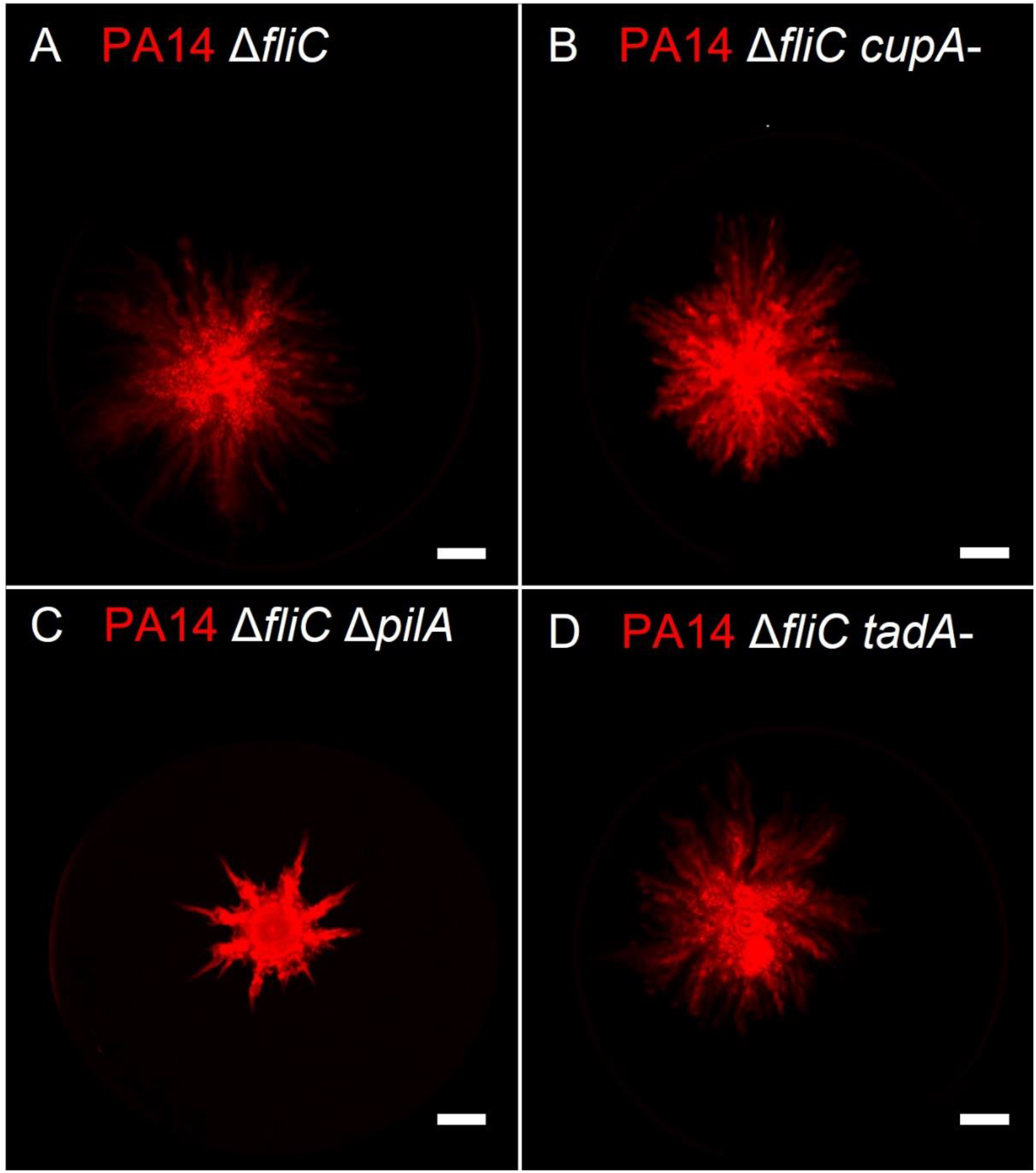
Co-swarming between *B. cenocepacia* K56-2 and *P. aeruginosa* PA14 aflagellated double mutants. Whole plate fluorescent scans of K56-2 with PA14 tagged with mCherry (red) were taken after overnight growth on semi-solid swarming M9DCAA medium. (**A)** Δ*fliC*, (**B)** Δ*fliC cupA*-, (**C)** Δ*fliC* Δ*pilA*, (**D)** Δ*fliC tadA*-. PA14 is tagged with mCherry (red). White scale bar = 1 cm.

**Figure 5.**
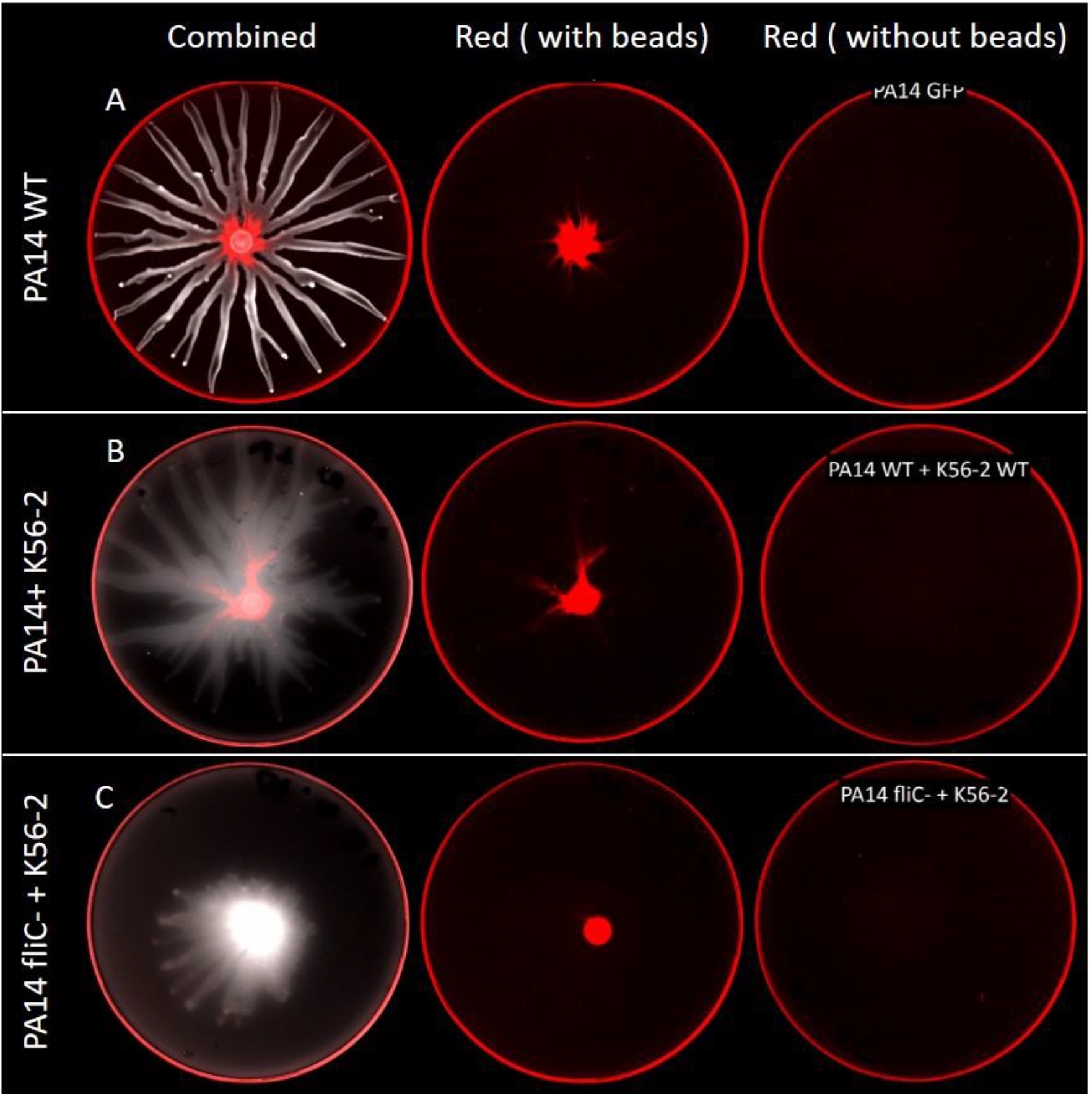
Localization of fluorescent polystyrene beads during co-swarming of *B. cenocepacia* K56-2 and *P. aeruginosa* PA14. An equal part of a suspension of 0.2 % red fluorescent 1 µm polystyrene beads were added to the bacterial suspension prior to inoculation (i.e., PA14:K56-2:Beads 1:1:1). Whole plate scans were taken after overnight growth on semi-solid M9DCAA. (**A)** PA14 alone, (**B)** PA14 with K56-2, (**C)** PA14 Δ*fliC* with K56-2.

## 3 Discussion

*P. aeruginosa* and *B. cenocepacia* are both environmental bacteria also recognized as opportunistic pathogens that can colonize the airways of people with cystic fibrosis. Both bacteria are motile and capable of swimming and swarming motilities. Under our experimental conditions for swarming assays, *P. aeruginosa* spreads across the surface of the agar gel to reach the border of the plate by forming a characteristic dendritic pattern; *B. cenocepacia* K56-2 is unable to spread in this fashion. Still, it can swarm on rich nutrient agar medium (25). *B. cenocepacia* possesses a polar flagellum and produces a yet-unidentified surfactant—both of which are required to swarm on agar. We found that K56-2 is unable to produce its surfactant on M9DCAA agar plates, resulting in a defect in swarming motility. We confirmed that lack of surfactant production is the limiting factor for swarming by supplementing exogenous rhamnolipids onto the surface of the agar gel and noting that it promoted swarming of *B. cenocepacia*. Accordingly, Bernier *et al*. (25) demonstrated that addition of spent supernatant of *P. aeruginosa* or of purified di-rhamnolipids into the medium allowed K56-2 to swarm on M9CAA. One striking difference here is the ability of K56-2 to form a dendritic swarm pattern under our conditions. This might be caused by the difference in the source of rhamnolipids used, but more likely by the mode of application: we added the rhamnolipids on the surface of the agar in the middle of the plate, producing a diffusion gradient, instead of being homogenously mixed into the medium prior to pouring into the plates. Rhamnolipids are responsible for the dendritic swarm pattern often displayed by *P. aeruginosa* (15, 31), and we know that this pattern requires this surfactant diffusion gradient (15). Our findings demonstrate that K56-2 is proficient at using rhamnolipids from PA14 to swarm.

Co-inoculating strains K56-2 and PA14 induced a remarkable modification of the swarm pattern compared to PA14 alone, translating into broader tendrils with a more uniform appearance, i.e. no presence of bulky tendril borders. In a study where a non-motile *E. coli* was added to a swarming *Acinetobacter baylyi*, the swarm pattern changed because of cell collision and the slowing down of the swarm front, leading to a flower-like pattern instead of the circular pattern of *A. baylyi* alone (32). Our microscopic observations suggest that the pattern deformation of the co-swarming between PA14 and K56-2 could result from *B. cenocepacia* segregation at the border of the tendril, causing a slowing-down of the swarm.

Under our conditions, an aflagellated PA14 (Δ*fliC*) mutant is incapable of swarming motility. However, it still retained the ability to move along in a swarm of *B. cenocepacia* K56-2 when co-inoculated. We hypothesized that *P. aeruginosa* translocates in the swarm by anchoring to K56-2 using one of its multiple cell appendages such as pili (type IVa, type IVb, tad) and cup fimbriae (CupA, CupB, CupC, CupD (33)). We tested double mutants harboring a *fliC* deletion combined with either the inactivation of CupA (*cupA3*-), Tad pilus (*tadA*-) or type IVa pilus (Δ*pilA*). None of these double mutants were hampered in their ability to co-swarm with K56-2. It could be that another appendage is used or that a combination participates in this interaction. We used fluorescent carboxylated-polystyrene beads to assess the ability of the swarm to carry around non-motile particles. Our reasoning was that beads could be either actively transported by the bacteria (lagging behind the swarm front) or passively pushed by the movement of the swarm (being pushed aside by the swarm front). Similar beads were shown to be displaced by *P. aeruginosa* during swimming motility (34). We observed that PA14 alone, and in co-swarming with K56-2, was able to move the beads through the tendrils. However, this displacement was abolished in the co-swarming with PA14 Δ*fliC* and K56-2. This suggests that transportation of the beads through the swarm cannot happen at the same time or that the flagellum acts as the adhesin for the beads, since it is known to be important for the initial attachment on polystyrene surfaces during biofilm formation (35). *P. aeruginosa* spread alongside *B. cenocepacia* could also be explained by *Burkholderia* triggering another type of motility from *P. aeruginosa* called sliding. Indeed, Murray and Kazmierczak reported a type of flagellum-independent motility called sliding that relies on rhamnolipids (36). They highlighted that this motility required the sensor kinase RetS. It would make sense that such motility could be triggered by K56-2 via a RetS-dependent pathway, since the presence of *Burkholderia thailandensis* was also reported to trigger the activity of RetS in *P. aeruginosa* (37). We thus explored this possibility, but co-swarming assays we conducted with Δ*fliC retS*-double mutants did not hinder the ability of PA14 to move in co-swarming (data not shown). Since the co-swarming of both K56-2 and PA14 Δ*fliC* requires rhamnolipids, it remains a possibility that PA14 simply follows the trail of K56-2 using this sliding motility mode. Unfortunately, very little is known about the mechanisms of sliding motility of *P. aeruginosa*.

There are some reports of motile bacteria carrying non-motile microorganisms as cargo. *Paenibacillus vortex* was reported to carry β-lactamase producing *E. coli* (38). In this case, *E. coli* serves as a shield against β-lactam antibiotics by continuously secreting beta-lactamases into the swarming medium. *P. vortex* also displays the ability to carry around conidia of *Aspergillus fumigatus* (39). This allows the conidia to spread and form mycelium which can further be exploited by *P. vortex* to cross gaps in the medium. *P. aeruginosa* was also shown to co-swarm with *Burkholderia cepacia*, which resulted in the ability the reach gentamicin-replete zones which were otherwise lethal (40). Our assays highlighted that *B. cenocepacia* K56-2 facilitates the spreading of an aflagellated mutant of *P. aeruginosa* PA14, while benefiting from rhamnolipids production by the latter.

*B. cenocepacia* and *P. aeruginosa* can both colonize the lung environment of cystic fibrosis patients (41). In lab growth assays, *P. aeruginosa* will often antagonize *B. cenocepacia* through the production of toxic effectors (e.g., hydrogen cyanide) and through competition for nutrients (mainly iron through pyoverdine) (26, 42, 43). However, they coexist in the swarming colony. Transcription of genes coding for some toxic effectors and competition factors such as hydrogen cyanide and siderophores is reduced in a swarming colony under our conditions (44, 45). This could contribute to explain why *B. cenocepacia* is able to thrive in the presence of *P. aeruginosa* when they swarm together. Interestingly, we did not observe a single species tendril breaking off in the co-swarming setting, although we know that K56-2 can spread by itself in the presence of rhamnolipids alone.

## 4 Conclusion

In this study, we highlight a peculiar interaction in multi-species swarming conditions where *P. aeruginosa* and *B. cenocepacia* can colonize a surface alongside each other. In this setting, cooperation seems to have precedence over competition: *P. aeruginosa* provides the surfactant needed by *B. cenocepacia* to spread across the surface whereas a non-motile mutant of *P. aeruginosa* is in return able to spread, helped by the presence of the motile *B. cenocepacia* cells. The mechanism by which this “hitchhiking” of *P. aeruginosa* occurs remains to be elucidated. This study provides new insights on another complex interaction between microbial species during social motility. The interactions between different bacterial species in a structured environment provides a condition favorable to interrogate bacterial social events.

## 5 Materials and methods

### 5.1 Strains, plasmids and growth conditions

Bacteria and plasmids used in this study are listed in **Table 1**. They are all derived from two parental strains, *P. aeruginosa* PA14 and *B. cenocepacia* K56-2 (46, 47). Bacteria were cultivated in Tryptic Soy Broth (TSB) (Difco) in a TC-7 roller drum (New Brunswick Scientific).

**Table 1:**
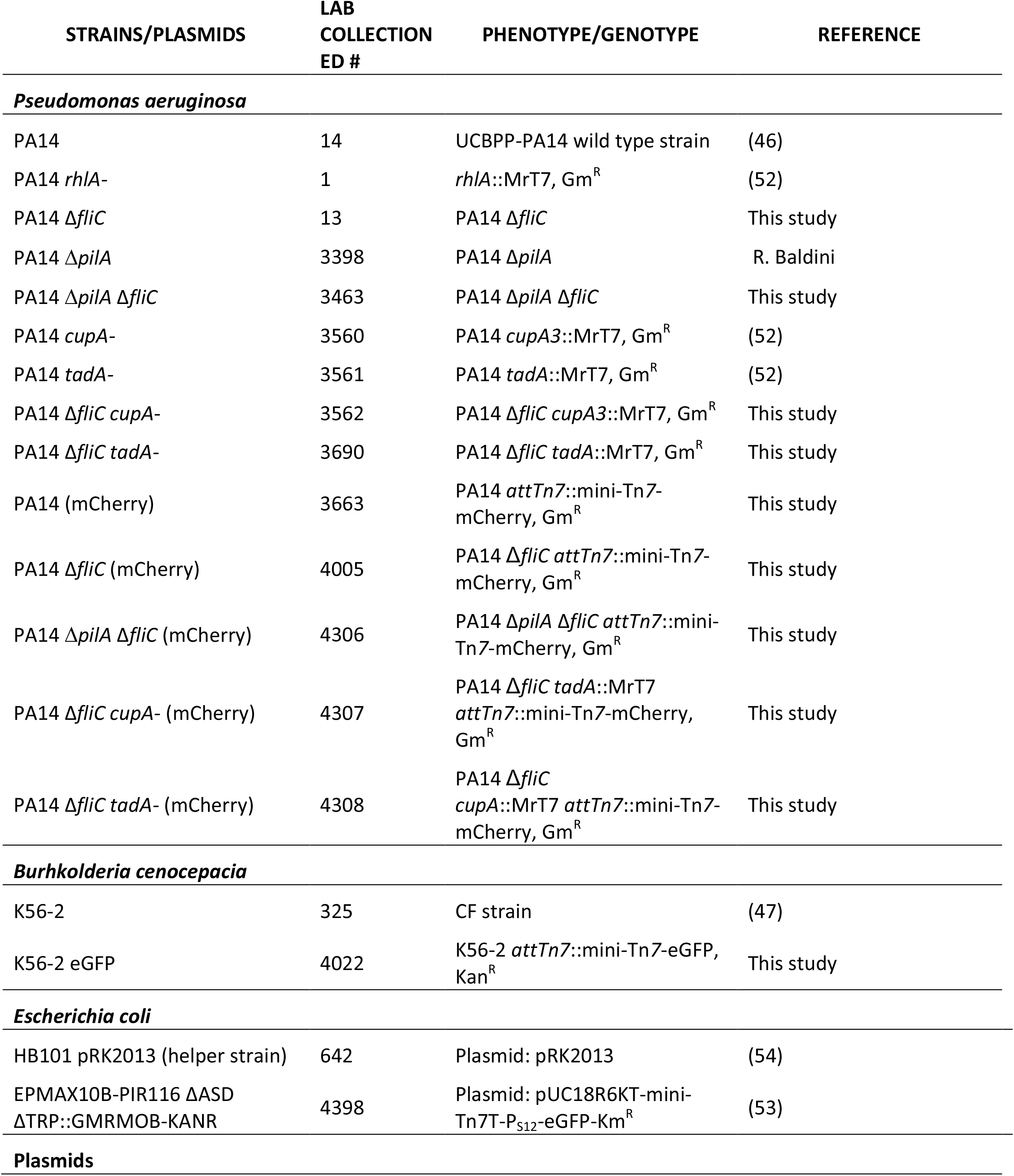
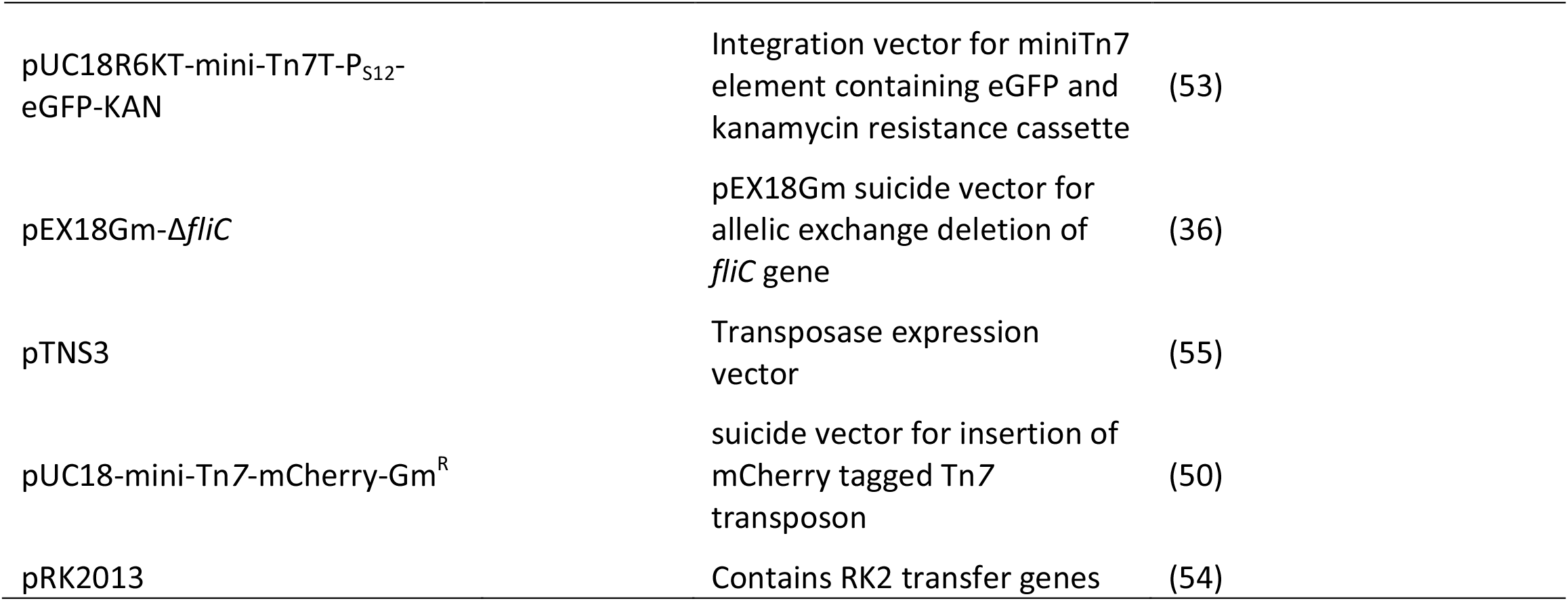
Strains/plasmids used in this study.

| STRAINS/PLASMIDS | LAB<br>COLLECTION<br>ED # | PHENOTYPE/GENOTYPE | REFERENCE |
| --- | --- | --- | --- |
| ***Pseudomonas aeruginosa*** |  |  |  |
| PA14 | 14 | UCBPP-PA14 wild type strain | (46) |
| PA14 *rhIA*- | 1 | *rhIA*::MrT7, Gm<sup>R</sup> | (52) |
| PA14 Δ*fliC* | 13 | PA14 Δ*fliC* | This study |
| PA14 Δ*pilA* | 3398 | PA14 Δ*pilA* | R. Baldini |
| PA14 Δ*pilA* Δ*fliC* | 3463 | PA14 Δ*pilA* Δ*fliC* | This study |
| PA14 *cupA*- | 3560 | PA14 *cupA3*::MrT7, Gm<sup>R</sup> | (52) |
| PA14 *tadA*- | 3561 | PA14 *tadA*::MrT7, Gm<sup>R</sup> | (52) |
| PA14 Δ*fliC* *cupA*- | 3562 | PA14 Δ*fliC* *cupA3*::MrT7, Gm<sup>R</sup> | This study |
| PA14 Δ*fliC* *tadA*- | 3690 | PA14 Δ*fliC* *tadA*::MrT7, Gm<sup>R</sup> | This study |
| PA14 (mCherry) | 3663 | PA14 *attTn7*::mini-Tn7-mCherry, Gm<sup>R</sup> | This study |
| PA14 Δ*fliC* (mCherry) | 4005 | PA14 Δ*fliC* *attTn7*::mini-Tn7-mCherry, Gm<sup>R</sup> | This study |
| PA14 Δ*pilA* Δ*fliC* (mCherry) | 4306 | PA14 Δ*pilA* Δ*fliC* *attTn7*::mini-Tn7-mCherry, Gm<sup>R</sup> | This study |
| PA14 Δ*fliC* *cupA*- (mCherry) | 4307 | PA14 Δ*fliC* *tadA*::MrT7<br>*attTn7*::mini-Tn7-mCherry,<br>Gm<sup>R</sup> | This study |
| PA14 Δ*fliC* *tadA*- (mCherry) | 4308 | PA14 Δ*fliC*<br>*cupA*::MrT7 *attTn7*::mini-Tn7-mCherry, Gm<sup>R</sup> | This study |
| ***Burkholderia cenocepacia*** |  |  |  |
| K56-2 | 325 | CF strain | (47) |
| K56-2 eGFP | 4022 | K56-2 *attTn7*::mini-Tn7-eGFP,<br>Kan<sup>R</sup> | This study |
| ***Escherichia coli*** |  |  |  |
| HB101 pRK2013 (helper strain) | 642 | Plasmid: pRK2013 | (54) |
| EPMAX10B-PIR116 ΔASD<br>ΔTRP::GMRMOB-KANR | 4398 | Plasmid: pUC18R6KT-mini-Tn7T-P<sub>S12</sub>-eGFP-Km<sup>R</sup> | (53) |
| **Plasmids** |  |  |  |

|  |  |  |
| --- | --- | --- |
| pUC18R6KT-mini-Tn7T-P<sub>S12</sub>-eGFP-KAN | Integration vector for miniTn7 element containing eGFP and kanamycin resistance cassette | (53) |
| pEX18Gm-Δ*fliC* | pEX18Gm suicide vector for allelic exchange deletion of *fliC* gene | (36) |
| pTNS3 | Transposase expression vector | (55) |
| pUC18-mini-Tn7-mCherry-Gm<sup>R</sup> | suicide vector for insertion of mCherry tagged Tn7 transposon | (50) |
| pRK2013 | Contains RK2 transfer genes | (54) |

### 5.2 Motility assays

Swarming motility assays were performed using M9DCAA semi-solid medium with an agar concentration of 0.5% (48). The medium was poured in 100 mm Petri dishes and dried for 25 minutes under laminar flow of a biosafety cabinet. Overnight bacterial cultures were washed twice with phosphate-buffered saline (PBS) and diluted to final OD_600_ of 3.0; OD_600_ was measured with a Nanodrop ND-1000 spectrophotometer (Thermo Fisher Scientific). For single-strain swarming, 5 μL of bacterial suspension was directly inoculated at the center of an agar plate. For dual-strain swarming, cells were mixed at a 1:1 ratio (based on OD_600_) and 5 μl of this mixed suspension was inoculated at the center of the plates. Plates were incubated overnight at 30°C or 37°C. Pictures were taken after incubation using a Panasonic ZS70 Lumix digital camera. At least three replicates were prepared for each assay.

### 5.3 Swarming with added rhamnolipids or plastic beads

A mix of purified *P. aeruginosa* rhamnolipids obtained from Jeneil Biosurfactant Co. (JBR-599, lot # 050629). This mixture containing 51 % monorhamnolipids, 45 % dirhamnolipids, and 3-hydroxy fatty acids was dissolved in MeOH at a concentration of 10,000 mg/L (49). A 5 μl drop was spotted onto the center of a swarming agar plate prepared as described above, with MeOH only used as a mock control. MeOH and rhamnolipids were left to dry for 10 minutes prior to inoculation with the bacterial suspension over the dried spot.

For assays including plastic beads, fluorescent 1 µm diameter carboxylated-polystyrene beads (FluoroSpheres, Invitrogen) were premixed with bacterial suspensions. The bead stock was washed with sterile water and diluted to a final concentration of 0.2 % (w/v). A volume of 5 µL of bead suspension was added to the bacterial suspension.

### 5.4 Generation of a Δ*pilA* Δ*fliC* double mutants

The *fliC* gene was deleted in a PA14 Δ*pilA* background by allelic exchange using vector pEX18Gm-Δ*fliC* as described in a previous study (16). Briefly, PA14 recipient cells received the vector via mating with an *E. coli* donor strain. First recombinants were selected on TSA with 15 µg/mL gentamicin, second recombinant were counter-selected on Tryptone Yeast agar with 10% sucrose. Potential Δ*fliC* were confirmed for their ability to swim in 0.3 % soft agar.

### 5.5 Fluorescence labelling of PA14 with mCherry

PA14 strains (wildtype, Δ*fliC*, Δ*pilA* Δ*fliC*) were labelled with mCherry at a single chromosomal site using pUC18T-miniTn*7*-P_A1/04/03_-*mCherry*-Gm^R^ (pBT277)(50). Both the Tn*7*-bearing vector and a transposase-expressing vector (pTNS3) were transferred into the recipient strains by electrotransformation (51). Cells that integrated the fluorescent marker were selected on TSA with 15 µg/mL gentamycin and confirmed for their ability to expression the mCherry fluorescent protein using a Typhoon FLA 9000 imaging system (GE Healthcare). The gentamicin resistance cassette was flipped by electrotransformation of the pFLP3a plasmid containing the FLP-recombinase into the newly produced fluorescent cells.

### 5.6 Inactivation of *cupA* and *tadA* genes in fluorescent cells

Further deletion of *cupA* and *tadA* genes was done with the transfer of transposon insertion from the PA14 transposon library (52). Genomic DNA from *cupA*::Mar2xT7 and *tadA*::Mar2xT7 mutants was extracted and transferred into the recipient into the recipient Δ*pilA* Δ*fliC attTn7*::mCherry background. Transformants were selected on TSA medium with 15 µg/mL gentamicin.

### 5.7 Fluorescent labelling of *B. cenocepacia* K56-2

Strain K56-2 was labelled with fluorescent protein eGFP from a Tn*7* delivery vector (53). The Tn*7*-bearing vector (pUC18R6KT-mini-Tn*7*T-P_S12_-eGFP-Km^R^) was transferred into the recipient cell via quad-mating between recipient (K56-2), donor *E. coli*, helper *E. coli* and transposase-bearing vector. Briefly, all four strains were grown overnight, diluted, and freshly grown for 4 hours at 37°C. A volume of 1 mL of cell each suspension was pelleted, washed and mixed before being spot inoculated onto a TSA plate. The suspension droplet was grown overnight; the resulting growth was spread onto TSA medium with 1,600 µg/mL kanamycin to select labelled K56-2. Acquired fluorescence was confirmed with using the Typhoon FLA 9000.

### 5.8 Fluorescent imaging of PA14 and K56-2 co-swarming

Fluorescent-labelled strains of PA14 and K56-2 were co-inoculated under swarming conditions as described above. Whole plates were then scanned using the Typhoon FLA 9000 with LPR filter and 532 nm wavelength laser for mCherry detection and BGP1 filter and 473 nm laser for green fluorescence, although subject to interference by *P. aeruginosa* autofluorescence. Close-up of tendrils were visualized with an Olympus Stereoscope using darkfield for the swarming colony and RFP filter for the fluorescence. Tendrils tips were visualized with a Zeiss LSM 780 confocal laser scanning microscope (CSLM). Agar pads bearing a tendril tip were carefully cut from the agar gel with a scalpel and placed onto a 3.5 mm coverslip-bottom dish. A section of 4 mm by 8 mm was imaged using a 20x objective in tile acquisition mode. Images were processed using Zeiss Zen Black Edition.

## 6 Acknowledgments

CM was recipient of Natural Sciences and Engineering Research Council of Canada (NSERC) Postgraduate Scholarships – Doctoral program. This research was funded by the NSERC under award number RGPIN-2020-06771.

We would like to thank Regina Baldini (Universidade de São Paulo) for providing us with a *pilA* deletion mutant of strain PA14. Special thanks to Salim Islam (INRS) for the help with stereomicroscopy, Jessy Tremblay for the help and troubleshooting with the confocal microscopy, and Nathalie Tufenkji (McGill University) for providing the fluorescent beads.

## 7 Author Contributions

Conceptualization, CM, ML and ED; methodology, CM and ML; investigation, CM; resources, ED; writing—original draft preparation, CM, ML and MCG; writing—review and editing, CM, MCG and ED; supervision, MCG and ED; project administration, ED; funding acquisition, ED. All authors have read and agreed to the published version of the manuscript.

